# Androgen exposure impairs neutrophil maturation and function within the infected kidney

**DOI:** 10.1101/2023.06.19.545598

**Authors:** Teri N. Hreha, Christina A. Collins, David A. Hunstad

**Author notes:** For correspondence: David A. Hunstad, MD, 660 S. Euclid Ave., MSC 8208-16-6, St. Louis, MO 63110, USA. Declaration of interests: D.A.H. serves on the Board of Directors of BioVersys AG, Basel, Switzerland, and has received research funding from BioAge Labs, Richmond, CA, USA. These activities are unrelated to the content of this manuscript.

## Abstract

Urinary tract infections (UTIs) in men are uncommon but carry increased risk for severe pyelonephritis and other complications. In models of *Escherichia coli* UTI, male C3H/HeN mice uniformly develop high-titer pyelonephritis (most with renal abscesses) in a testosterone-dependent manner, but the mechanisms underlying this phenotype are unknown. Here, using female mouse models, we show that androgen exposure impairs neutrophil maturation in the upper and lower urinary tract, compounded by an additional reduction of neutrophil function specifically within the infected kidney, enabling persistent high-titer infection and promoting abscess formation. Following intravesical inoculation with uropathogenic *E. coli* (UPEC), kidneys of androgen-exposed C3H mice showed delayed local pro-inflammatory cytokine responses while robustly recruiting neutrophils. These were enriched for an end-organ-specific population of aged but immature neutrophils (CD49d+, CD101–). Compared to their mature counterparts, these aged immature kidney neutrophils exhibited reduced functions *in vitro*, including impaired degranulation and diminished phagocytic activity, while splenic, bone marrow, and bladder neutrophils did not display these alterations. Further, aged immature neutrophils exhibited little phagocytic activity within intratubular UPEC communities *in vivo*. Experiments with B6 conditional androgen receptor (AR)-deficient mice indicated rescue of the maturation defect when AR was deleted in myeloid cells. We conclude that the recognized enhancement of UTI severity by androgens reflects urinary tract-specific impairment of neutrophil maturation (largely via cell-intrinsic AR signaling) and kidney-specific reduction in neutrophil antimicrobial capacity, resulting in failure to control renal bacterial infection.

## Introduction

Urinary tract infections (UTIs) are extremely prevalent, and over 80% are caused by uropathogenic *Escherichia coli* (UPEC). Although the majority of UTIs occur in females, the incidence of UTI in males is higher than in females among infant and elderly populations (1–6). Male UTI is also more complicated than in females, with renal infections carrying increased morbidity and mortality and significantly increasing the risk for hypertension, renal scarring, and chronic kidney disease (1,7–15).

Recent work in preclinical models has demonstrated that this increased propensity for severe UTI in males is testosterone dependent. Female C57BL/6 and C3H/HeN mice exhibit a more robust cytokine response in their bladders compared to male mice within 24 hours post infection (hpi) (16,17). This early response results in increased early recruitment of immune cells, including neutrophils, monocytes and macrophages, favoring resolution of the UTI in females (17). In C3H/HeN mice, which feature vesicoureteral reflux (a major risk factor for upper-tract UTI in children (18–20)), male mice develop chronic cystitis and pyelonephritis marked by renal abscess formation, while normal females resolve infection generally within 7 days (16,21). Castrated or androgen receptor (AR)-deficient males are able to successfully resolve infection, while androgenization of female mice recapitulates the severe UTI susceptibility phenotype, with organ bacterial loads and abscess prevalence comparable to those of male mice (16,17,21). While these abscesses harbor large numbers of neutrophils, there is no discernible reduction in kidney bacterial load in androgen-exposed C3H/HeN and C57BL/6 mice through at least 28 days post infection (dpi), indicating that the neutrophil response to pyelonephritis in the androgenized host is insufficient to control or eradicate infection (16,17,22).

Testosterone is generally considered to be immunosuppressive in a variety of diseases, dampening pro-inflammatory mediators such as TNFα and iNOS while increasing synthesis of anti-inflammatory mediators such as IL-10 and TGFβ (23–26). Males are known to have higher circulating neutrophil counts than females (27,28). Women with polycystic ovary syndrome (PCOS; a well-recognized hyperandrogenic state) have higher circulating neutrophil counts than healthy females, and these numbers are normalized in PCOS patients receiving combination treatment that includes the AR antagonist flutamide (29). Mice with germline deletion of *Ar* have significantly lower circulating neutrophil counts (30).

Both male and female neutrophils and their precursors express high levels of AR (31), and testosterone exposure has been shown to affect neutrophil function. In *in vitro* assays, male neutrophils exhibited reduced chemotactic and phagocytic capacity (32,33). After 12 weeks of testosterone therapy, peripheral neutrophils from transgender men showed higher IL-6 and TNFα production while also exhibiting increased adherence and reduced rolling velocity on endothelial cells (34). In animal models, testosterone is associated with a suppressed innate immune response to acute bacterial cystitis, particularly through reduction of IL-17 and γδ T cells (17).

Similarly, androgens were shown to enhance recruitment of neutrophils, but also to stimulate their IL-10 production and inhibit their phagocytic ability, in acute bacterial prostatitis (35). In noninfectious renal ischemia/reperfusion injury, males exhibited a stark increase in recruitment of bone marrow-derived neutrophils to the kidney compared to females, reflecting increased β2 integrin expression driven by sex-specific effects of CXCL5 (36).

Neutrophil maturation normally occurs in the bone marrow, with immature neutrophils (CD101– in mice, CD10– in humans) exhibiting incomplete nuclear development and reduced granular content. Immature neutrophils are prematurely released from the bone marrow in response to high levels of cytokines such as G-CSF and GM-CSF and can migrate to sites of infection at the same rate as mature neutrophils (37,38). Immature neutrophils are immunostimulatory, secreting pro-inflammatory cytokines such as type I interferons, TNFα, and IFNγ, and their accumulation at inflammatory sites correlates with disease progression (38–40). While immature and mature neutrophils exhibit comparable chemotaxis toward inflammatory stimuli, immature neutrophils exhibit reduced granular function (as measured by myeloperoxidase activity) and phagocytic capacity *in vitro* (41,42).

Murine neutrophils typically exhibit a half-life of 6-12 h in circulation (43). However, some neutrophils may remain in circulation or in tissues longer, and these aged neutrophils may act as a first line of defense in organ inflammation. Aged neutrophils that have circulated in mice for at least 48 h express low levels of CD62L, and high levels of CXCR4 and the integrin subunits CD11b and CD49d, making them better able to adhere to tissue endothelia (44–46). In states of inflammation (e.g., LPS exposure), aged neutrophils traffic to the inflammatory site faster, produce higher levels of IL-6, and are better able to phagocytose bacteria than younger neutrophils (46). However, accumulation of aged neutrophils in tissues can promote further inflammation and tissue damage (47–49).

Here, we demonstrate that neutrophil-rich renal abscess formation during UTI in androgenized C3H/HeN mice correlates with accumulation of a distinct tissue-specific population of neutrophils that are aged but remain immature. Although these aged immature neutrophils properly traffic to foci of bacterial infection in the kidney, they exhibit reduced degranulation and phagocytic capacity, rendering them less effective at controlling infection. We further demonstrate that the maturation failure of neutrophils within the kidney is largely attributable to cell-intrinsic AR signaling. Our findings reveal a key cellular mechanism by which androgen exposure predisposes to severe pyelonephritis and renal abscess formation.

## Results

### Persistent, high-titer pyelonephritis in androgenized mice is characterized by continuous neutrophil recruitment

As outlined above, testosterone underlies the enhanced severity of UTI we have previously observed in male mice. To mechanistically investigate the effect of testosterone on the neutrophil response to pyelonephritis, we exposed C3H/HeN females to androgen via injection of testosterone cypionate, yielding serum testosterone levels approximating the biological range of adult C3H/HeN males (50), prior to UPEC inoculation.

Concordant with our prior results in male and testosterone pellet-implanted female C3H/HeN mice (16,21), and in testosterone cypionate-treated C57BL/6 females (22,51), androgenized C3H/HeN females developed chronic cystitis (**Figure 1A**) and pyelonephritis (**Figure 1B**), with consistently high bacterial loads across all measured time points. Androgen exposure resulted in increased CD45+ cells in the kidney (as a percentage of live cells, and in absolute number) prior to initiation of experimental UTI (i.e., in naïve mice), and CD45+ cell populations increased within the kidneys of androgenized mice as infection progressed (**Figure 1C, E**). By 14 dpi, androgenized mice had significantly more CD45+ cells in the kidneys than at 1 dpi (p = 0.0027), while vehicle-treated females had significantly fewer than at 1 dpi (p = 0.00067; **Figure 1E**). By 10 dpi, neutrophils made up ∼60% of the CD45+ population in the kidney in androgenized mice, significantly more than in vehicle-treated mice (**Figure 1D, F**). This influx of neutrophils to the kidney occurred before any measurable increase in neutrophils in the peripheral blood (as a proportion of CD45+ cells; **Figure S1A**), indicating that neutrophils released from bone marrow were homing to the site of infection. Indeed, vehicle-treated and androgenized mice exhibited similar neutrophil counts in peripheral blood 10 dpi (**Figure S1B**). While neutrophil recruitment to the kidney in vehicle-treated mice peaked 1 dpi, this process continued in androgenized mice, with significantly more neutrophils present 14 dpi than at 1 dpi (p = 0.0027) (**Figure 1F**). Among other CD45+ cell types, only T cells significantly increased between 1 and 14 dpi in the kidneys of androgenized mice (**Figure S1C-H**).

**Figure 1.**
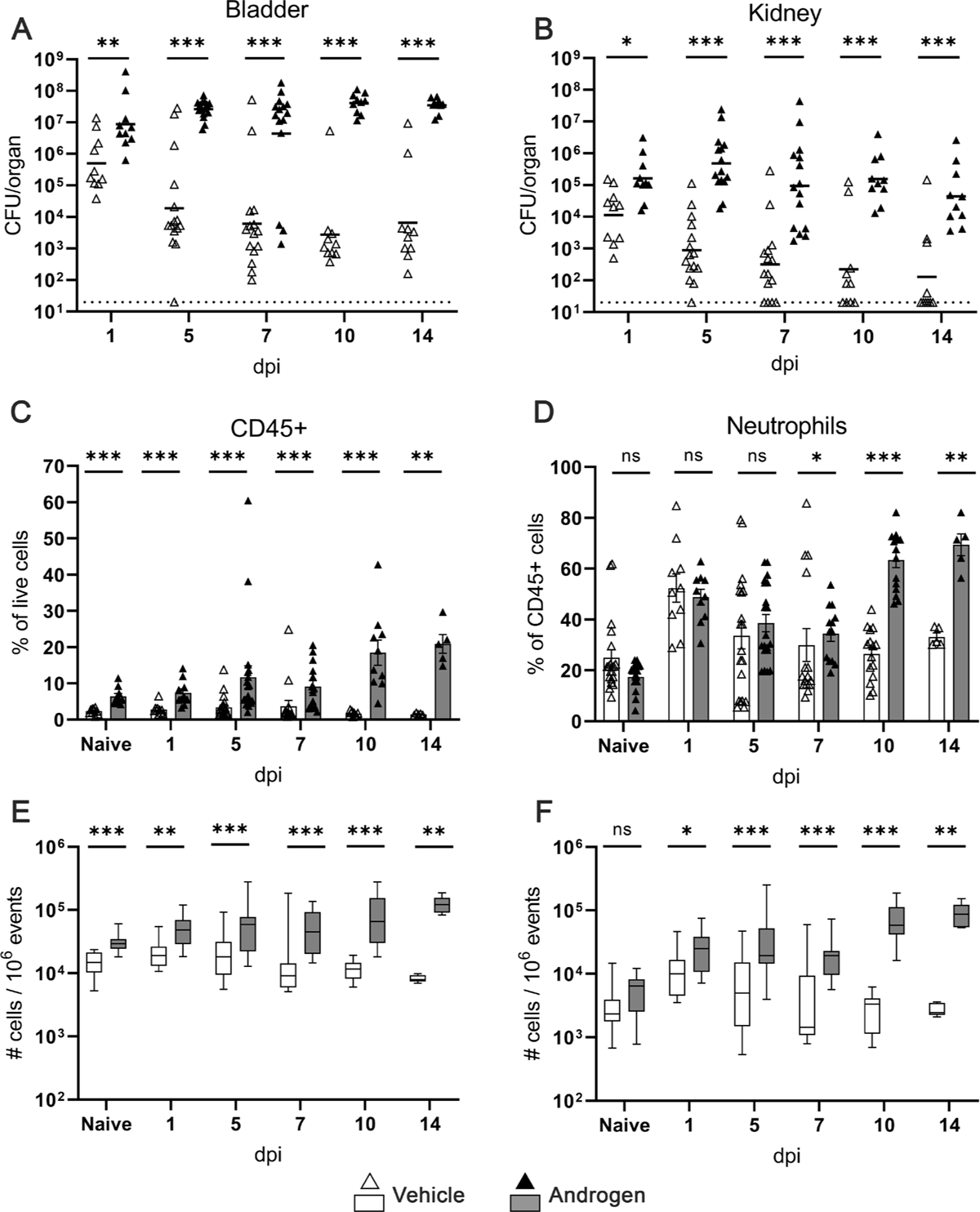
Androgen exposure enables chronic UTI with continuous neutrophil recruitment. (A-B) Timeline of bladder and kidney bacterial loads after UPEC inoculation in vehicle-treated (open triangles) and androgenized (filled triangles) C3H/HeN mice. Lines indicate geometric mean. (C) CD45+ cell recruitment to the kidney over time as a percentage of live cells, by flow cytometry. (D) Neutrophil (CD45+, Ly6G+) recruitment to the kidney over time as a percentage of CD45+ cells. (E-F) Timeline of the number of CD45+ cells (E) and neutrophils (F) in the kidneys throughout the course of infection, expressed per 1 million events, in vehicle-treated (white boxes) or androgenized mice (gray boxes). Bars indicate mean with SEM. Each symbol represents a single mouse; n = 5-15 per condition. *p < 0.05, **p < 0.01, ***p < 0.001 by Mann-Whitney U test. CFU, colony-forming units. ns, not significant.

### Cytokine responses to UTI are delayed in the androgenized kidney

We previously reported that male and androgenized female mice exhibit elevated expression of pro-fibrotic and pro-inflammatory mediators, “priming” the mouse for an aberrant response to UTI (16,51). This effect was recapitulated in the kidney cytokine profile of androgenized C3H/HeN females, which featured significantly higher levels of G-CSF, IL-1α, IL-1β, and IL-6 than vehicle-treated females prior to UTI (naïve; **Figure 2**). Following initiation of UTI, there were no increases in these cytokines, or in IL-17 or KC (CXCL1), measured in vehicle-treated mice (**Figure 2**), likely reflecting that whole-kidney cytokine analysis is insensitive to changes associated with a modest and localized infection. In androgenized mice 7 dpi, whole-kidney levels of the neutrophil-recruiting cytokines IL-1α, IL-1β, and IL-6, were unchanged from baseline (day 0; p = 0.9999, 0.5476, 0.9999, respectively), while IL-17, G-CSF, and KC (CXCL1) were significantly increased (**Figure 2**; p = 0.0079, 0.0159, 0.0159, respectively); notably, at this time point renal abscess is already established (21). Whole-kidney levels of other neutrophil-recruiting (IL-3, GM-CSF) or inhibitory cytokines (IL-4, IL-10) were not significantly altered at measured time points after infection (data not shown).

**Figure 2.**
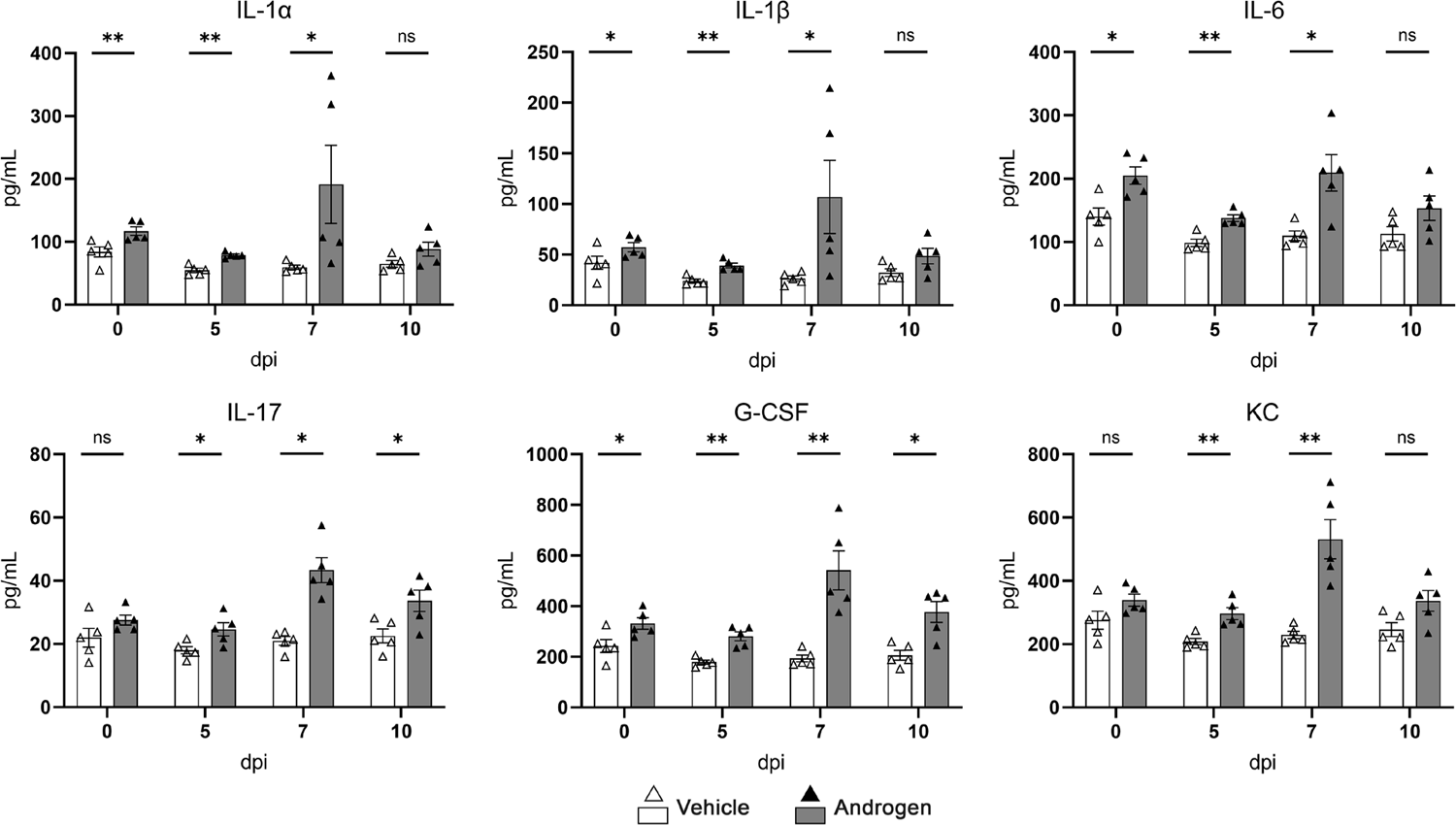
Local production of neutrophil recruiting cytokines in the kidney is delayed in androgenized mice. Timeline of whole-kidney levels of the indicated cytokines in vehicle-treated (white bars, open triangles) and androgenized mice (gray bars, filled triangles). Bars indicate mean with SEM. Each symbol represents a single mouse; n = 5 per time point. *p < 0.05, **p < 0.01 by Mann-Whitney U test. ns, not significant.

### Androgenized mice harbor a distinctly large population of aged, immature neutrophils in infected kidneys

To investigate how high-titer pyelonephritis persists in the androgenized kidney despite robust neutrophil recruitment, we next interrogated the age and maturity of recruited neutrophils. Using selected flow cytometric markers for age (CD49d) and maturity (CD101), we found that most of the neutrophils (CD45+, Ly6G+) in the kidneys or peripheral blood of vehicle-treated mice were either young and immature (CD49d–, CD101–) or aged and mature (CD49d+, CD101+; **Figure 3A,C**). While neutrophils in the peripheral blood of androgenized mice aged and matured similarly to those of vehicle-treated mice (**Figure 3D**), the kidneys of androgenized mice accumulated a sizable population of neutrophils that were aged but immature (CD49d+, CD101–; **Figure 3B**). Analysis of additional neutrophil markers demonstrated that aged or mature neutrophils were more likely to be CD62L^lo^, CXCR2^lo^, CXCR4+, CD11b^hi^, and young or immature neutrophils were CD62L^hi^, CXCR2^hi^, CXCR4-, CD11b^lo^ (**Figure S2I-L**). As expected, the aged immature population had intermediate expression of all of these markers, compared to populations that were strictly gated on either age or maturity. Ly6G+, CD101–, CD49d+ neutrophils have been previously described in the bone marrow as committed neutrophil precursors (37); however, in our model, these cells were found in the kidney and not in peripheral blood, indicating they are tissue-infiltrated aged neutrophils rather than neutrophil precursors released prematurely from bone marrow. Vehicle-treated mice exhibited expansion of multiple neutrophil age/maturity subtypes 1 dpi, dominated by swift recruitment of aged mature neutrophils, with a return to baseline by 10 dpi. In contrast, in androgenized mice, expansion of young immature and aged (both mature and immature) neutrophils persisted through 14 dpi (**Figure S2E-H**). Viewed another way, the proportion of aged immature neutrophils remained low as infection resolved in vehicle-treated mice (**Figure 3E and S2C,G**) but had risen significantly in androgenized mice by 10 and 14 dpi (p < 0.01; **Figure 3F and S2C,G**). In addition, the proportion of aged mature neutrophils in the kidneys, though equal at baseline, was significantly lower in androgenized mice than in vehicle-treated mice early in infection (1 and 5 dpi; p < 0.05), while the proportion of young immature neutrophils was significantly increased in androgenized mice at these time points (p < 0.01; **Figure 3E,F and S2A,D**). Further, the proportion of aged mature neutrophils in the kidneys at later time points (10 dpi and 14 dpi) were significantly reduced in androgenized mice (p < 0.05 or p < 0.01; **Figure 3E,F and S2B**). These data indicate a failure of neutrophil maturation at both acute and chronic time points within the infected kidney in androgenized hosts.

**Figure 3.**
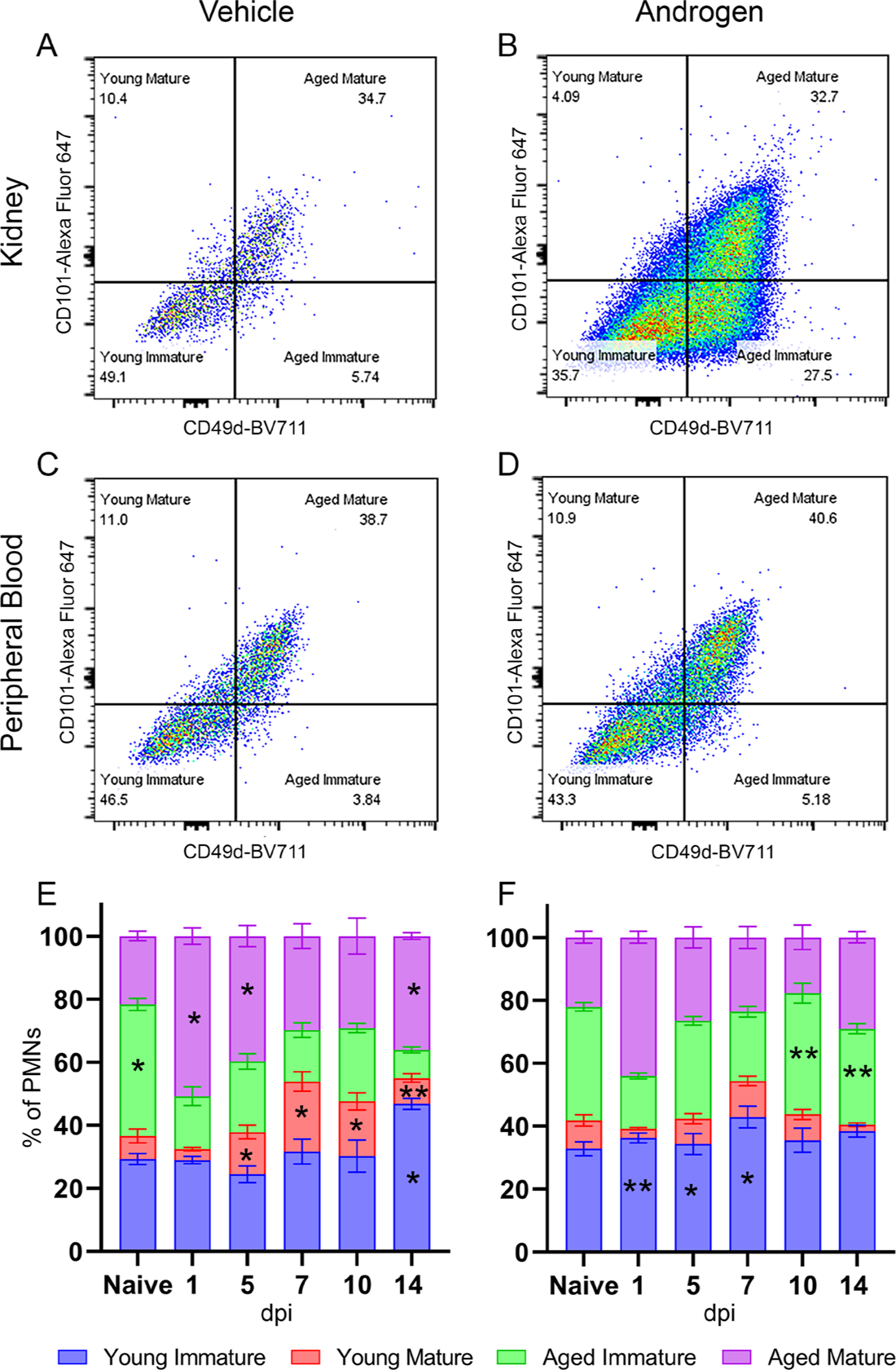
UPEC-infected kidneys in androgenized mice harbor a distinctive population of aged immature neutrophils. Representative gating of neutrophils (CD45+, Ly6G+) for age (CD49d) and maturity (CD101) in kidneys (A-B) or peripheral blood (C-D) from vehicle-treated and androgenized mice 14 dpi. (E-F) Timeline of the proportion of neutrophils of each subtype (young immature [blue bars], young mature [red], aged immature [green], aged mature [purple]) throughout the course of infection in vehicle-treated and androgenized mice. Bars indicate mean with SEM. n = 5-15 per condition and time point. *p < 0.05, **p < 0.01 by Mann-Whitney U test.

### Maturation impairment is driven by androgen receptor signaling on myeloid cells

To determine if the androgenic impairment of neutrophil maturation was due to cell type-specific androgen receptor signaling, we generated mice lacking expression of AR in myeloid cells (including neutrophils; LysM-Cre × AR^f/f^) or in renal epithelium (Ksp-Cre × AR^f/f^). Of note, the genetically modified parent strains exist in the C57BL/6 background, which does not feature the vesicoureteral reflux characteristic of C3H/HeN mice; we previously demonstrated that androgenized C57BL/6 females maintain lower kidney bacterial loads than similarly inoculated C3H/HeN mice but are susceptible to chronic pyelonephritis and renal scarring (22,51). Infection of androgenized Ksp-Cre × AR^f/f^ and LysM-Cre × AR^f/f^ mice resulted in high-titer cystitis and pyelonephritis 7 dpi, similar to that of androgenized Cre^-^AR^f/f^ littermate controls (**Figure S3A**), accompanied by a similar influx of neutrophils to the kidney, both as a percentage of CD45+ cells and in absolute number (**Figure S3B, C**).

Androgenization of Cre^-^AR^f/f^ control mice resulted in an increase in the proportion of immature neutrophils in the kidney 7 dpi compared to vehicle-treated Cre^-^AR^f/f^ littermate controls (white vs gray bars, **Figure 4A**), recapitulating our observations in C3H/HeN mice. This effect was incompletely but significantly mitigated in androgenized LysM-Cre × AR^f/f^ mice (gray vs pink bars, **Figure 4A**); furthermore, compared to androgenized Cre^-^AR^f/f^ control mice, the kidneys of infected LysM-Cre × AR^f/f^ mice displayed a significant decrease in aged immature neutrophils (**Figure 4B**). These effects were not seen in Ksp-Cre × AR^f/f^ mice (gray vs green bars, **Figure 4A,B**). Correspondingly, neutrophil maturation in the kidney was rescued in these LysM-Cre × AR^f/f^ mice, matching vehicle-treated Cre^-^AR^f/f^ littermates (**Figure 4C,D,F**), while maturation was not rescued in Ksp-Cre × AR^f/f^ mice (**Figure 4D,E**). Taken together, these data demonstrate that kidney neutrophil maturation was similarly impaired by androgen exposure in both B6 and C3H models, and that the androgen-dependent neutrophil maturation defect is driven substantially by myeloid cell-specific AR signaling. Of note, neutrophil maturity in the kidneys of B6 mice did not correlate with neutrophil age as directly as was seen in C3H/HeN mice.

**Figure 4.**
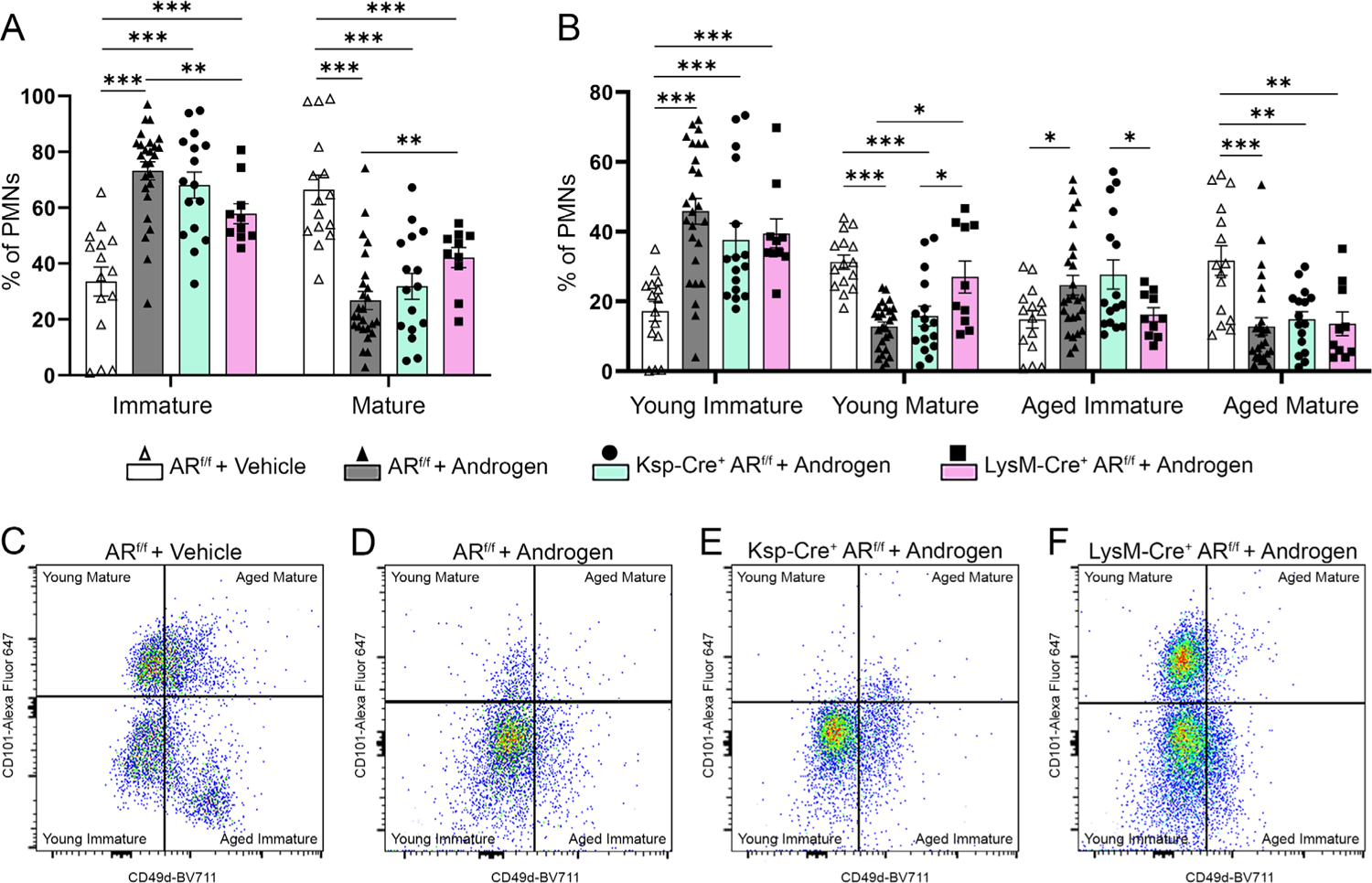
Inhibition of AR signaling in neutrophils substantially restores their maturation in the infected kidney. (A) Proportion of neutrophils that were immature (CD101–) or mature (CD101+) 7 dpi in vehicle-treated Cre^-^ AR^f/f^ (open triangles, white bars), androgenized Cre^-^ AR^f/f^ (closed triangles, gray bars), androgenized Ksp-Cre × AR^f/f^ (squares, green bars), or androgenized LysM-Cre × AR^f/f^ (circles, pink bars) C57BL/6 mice. (B) Proportion of neutrophils that were young immature, young mature, aged immature, and aged mature in vehicle-treated Cre^-^AR^f/f^, androgenized Cre^-^AR^f/f^, androgenized Ksp-Cre × AR^f/f^, or androgenized LysM-Cre × AR^f/f^ C57BL/6 mice 7 dpi (symbols as in panel A). Bars indicate mean with SEM. (C-F) Representative pseudocolor plots of neutrophil age (CD49d) and maturity (CD101) by flow cytometry in vehicle-treated Cre^-^AR^f/f^, androgenized Cre^-^AR^f/f^, androgenized Ksp-Cre × AR^f/f^, or androgenized LysM-Cre × AR^f/f^ C57BL/6 mice. n = 15-26 per mouse strain. *p < 0.05, **p < 0.01, ***p < 0.001 by Mann-Whitney U test. Comparisons with p-values > 0.05 are not indicated.

Specifically, while C3H/HeN mice harbored very few young mature neutrophils in the kidneys 7 dpi (**Figure 3A,B**), most of the neutrophils in B6 kidneys 7 dpi were young, with few aged neutrophils present in the kidneys of any group (**Figure 4C-F**).

### Phagocytosis of UPEC is inhibited in neutrophils of androgenized mice

To further interrogate the effect of androgen exposure on neutrophil functions, leukocytes were isolated from the spleen, bone marrow, or kidneys of vehicle-treated or androgenized mice that had no prior infection (naïve) or were 7 days post UPEC infection. These isolated cells were tested *ex vivo* for their capacity to phagocytose GFP-expressing UPEC. Ly6G+ neutrophils harvested from any sampled site in vehicle-treated mice had similar phagocytic capacity, with ∼25-40% positivity for GFP (white bars, **Figure 5A**). Androgen exposure did not affect phagocytosis by neutrophils isolated from either spleen or bone marrow (**Figure 5A**). However, kidney neutrophils from naïve androgenized mice showed significantly less phagocytic activity than their vehicle-treated counterparts (∼20% GFP+, p < 0.01); this phenotype was even more pronounced in kidney neutrophils harvested from androgenized mice 7 dpi (∼10% GFP+, p = 0.007 vs vehicle) (**Figure 5A**). Of note, these differences were not due to limitation of CFU available for engulfment at the end of incubation in androgenized conditions (**Figure S4A**).

**Figure 5.**
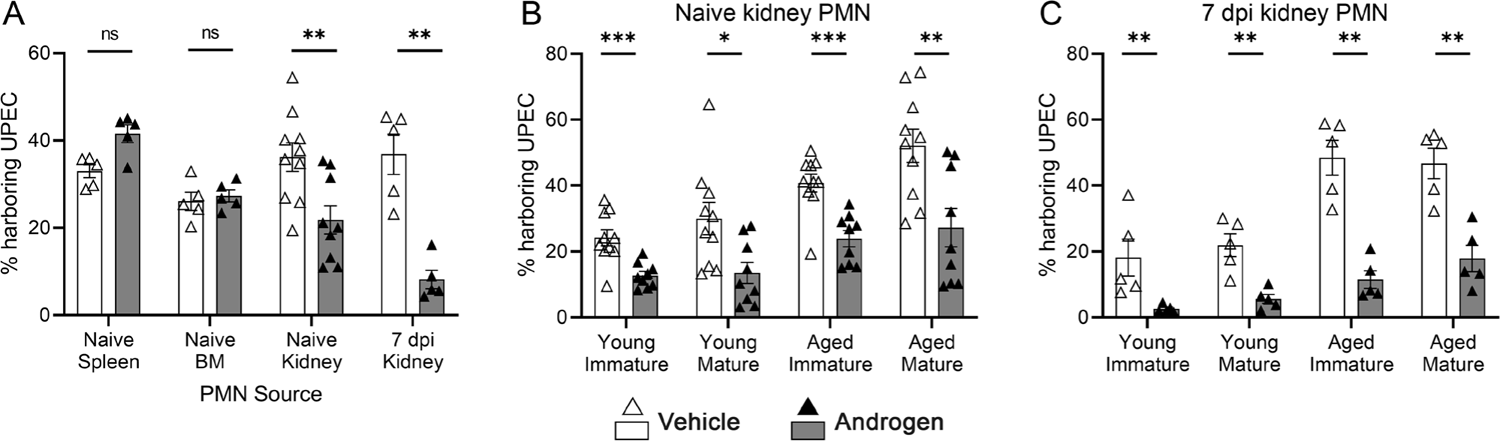
Kidney neutrophils in androgenized mice exhibit reduced phagocytic capacity. (A) Percentage of neutrophils that phagocytosed GFP+ UPEC after isolation from the spleen, bone marrow, or kidney in the naïve state, or the kidney 7 dpi, in vehicle-treated (open triangles) or androgenized mice (filled triangles). (B-C) Percentage of young immature, young mature, aged immature, and aged mature neutrophils isolated from naïve or 7-dpi kidneys in vehicle-treated or androgenized mice. Bars indicate mean with SEM. Each symbol represents a single mouse; n = 5-10 per group. *p < 0.05, **p < 0.01, ***p < 0.001 by Mann-Whitney U test. ns, not significant.

Reduced phagocytic capacity associated with androgen exposure was also observed in other (CD45+ Ly6G–) leukocyte populations from the kidney, but not from the spleen (**Figure S4B**), indicating that the local environment of the androgen-exposed kidney limits the functional potential of phagocytes, even in the uninfected state.

We next quantified phagocytic capacity of kidney neutrophils according to age/maturity subtypes. Among neutrophils from the kidneys of naïve mice, aged neutrophils exhibited more robust phagocytosis of UPEC than young neutrophils, and mature neutrophils were slightly more effective than their immature counterparts, consistent with prior reports (41,42,46), in both vehicle-treated and androgenized groups (**Figure 5B**). Furthermore, phagocytic capacity was significantly higher in neutrophils (across all subtypes) from vehicle-treated mice than from androgenized mice (**Figure 5B**). These androgen-dependent differences across all subtypes were even more striking among neutrophils from the kidneys of mice infected with UPEC for 7 days (**Figure 5C**). Moreover, phagocytic capacity of immature (both young and aged) kidney neutrophils diminished significantly in androgenized mice between the naïve and 7-dpi state (gray bars in **Figure 5B vs 5C**, p = 0.0009 and 0.007 respectively), while there were no differences in phagocytic function of any neutrophil subtype in vehicle-treated mice between the naïve and 7-dpi state (white bars in **Figure 5B vs 5C**).

### Aged immature neutrophils accomplish little phagocytosis in the UPEC intratubular niche

We previously showed that intratubular kidney bacterial communities (KBCs) within the developing renal abscess are surrounded by a large population of Ly6G+ neutrophils (16). Here, we localized the kidney neutrophils by age and maturity subtypes in relation to the KBC by immunofluorescence microscopy. As shown before, KBC-bearing tubules were surrounded by Ly6G+ neutrophils; most of these stained as young and immature (**Figure 6A**). Neutrophils of each subtype could be found within the KBC itself (**Figure 6B**), suggesting that functional defects in aged immature neutrophils do not include outright failure of trafficking to intratubular UPEC. However, *E. coli* staining co-localized exclusively with mature neutrophils (CD101+), with no *E. coli* positivity in aged immature neutrophils (**Figure 6B**). These data indicate that while infiltrating neutrophils (regardless of maturation status) are able to reach UPEC within KBCs, the aged immature neutrophils that are much more prominently represented in the androgenized host exert little to no phagocytic activity in that niche.

**Figure 6.**
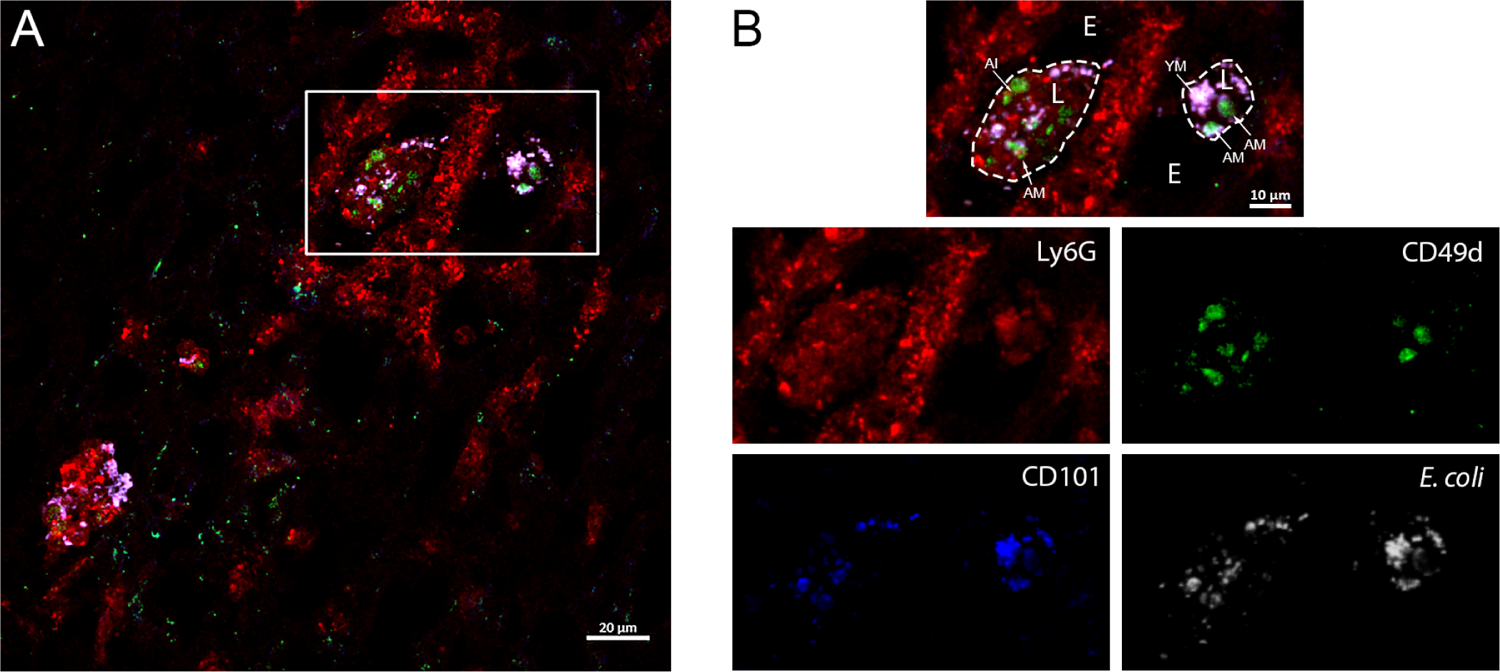
Only mature neutrophils exhibit phagocytic function within kidney bacterial communities. Representative immunofluorescence localization of neutrophil subtypes near and within KBCs of an androgenized mouse 10 dpi. White, *E. coli*; red, Ly6G (neutrophils); green, CD49d (age); blue, CD101 (maturity). Inset in (A) magnified in the panels in (B). E, epithelium; L, tubular lumen. Arrows indicate different neutrophil subtypes: YM, young mature; AI, aged immature; AM, aged mature.

### Degranulation by kidney neutrophils in response to UTI is blunted in androgenized mice

Neutrophils bear multiple types of granules with specific antimicrobial contents, released in a defined order – secretory vesicles, followed by tertiary (gelatinase) granules, then secondary (specific) granules, and finally primary (azurophilic) granules (52–54). Returning to the C3H/HeN model, we next assessed the extent and tempo of degranulation in kidney neutrophils, using flow cytometry markers detectable on the cell surface after selected granules fuse with the cell membrane. The release of secretory vesicles and primary granules from kidney neutrophils of vehicle-treated mice rose 1-7 dpi (compared to the naïve state; **Figure 7A,F**), then fell to baseline by 10 dpi in concert with resolving bacterial loads (see **Figure 1A,B**). In contrast to vehicle-treated mice, degranulation by kidney neutrophils from androgenized mice was blunted 1 dpi and remained reduced through 7 dpi (**Figure 7A,F**). This suppressed functional response in kidney neutrophils of the androgenized host correlates temporally with failure to control bacterial loads in the tissue (**Figure 1A,B**) and formation of abscesses (21). Following this ineffective initial response, release of primary granules increased 10 dpi, toward levels observed days earlier in vehicle-treated mice (**Figure 7F**). This degranulation 10 dpi was specifically attributable to the mature neutrophil populations (**Figure 7H,J**), consistent with previously published literature showing that immature neutrophils have reduced granular function (41,52). Indeed, aged immature neutrophils exhibited significantly less degranulation activity 10 dpi compared to their aged mature counterparts in both vehicle-treated and androgenized mice (p = 0.055 for secretory vesicles, 0.008 for primary granules in vehicle-treated mice; p = 0.0317 for secretory vesicles, 0.008 for primary granules in androgenized mice). As observed earlier, these aged immature neutrophils comprise nearly 40% of the kidney neutrophil population 10 dpi in androgenized mice (p<0.01 vs vehicle-treated; **Figure 3F and S2C**), likely compounding the inability of androgenized mice to effectively clear UPEC from the kidneys.

**Figure 7.**
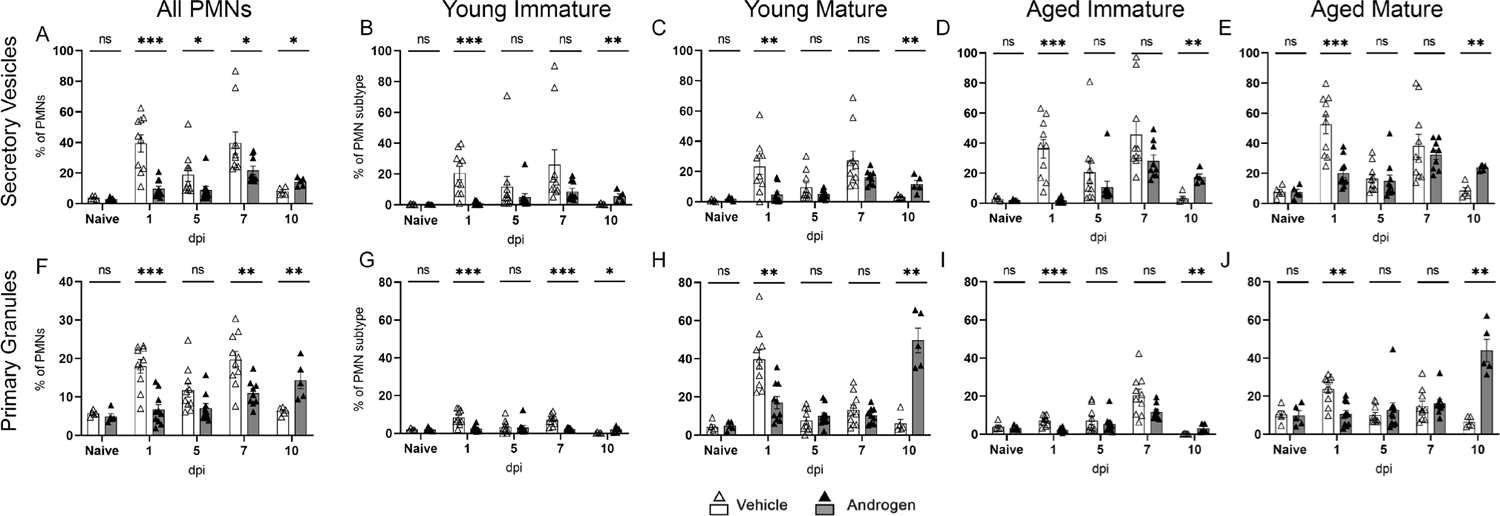
Degranulation by kidney neutrophils is blunted and delayed in androgenized mice. Release of secretory vesicles (CD18+ CD11b+; A-E) and primary granules (CD63+; F-J), expressed as a proportion of all neutrophils (A, F) or of each neutrophil subtype at the indicated time points post UPEC infection in vehicle-treated (open triangles) or androgenized mice (filled triangles). Bars indicate mean with SEM. Each symbol represents a single mouse; n = 5-10 per time point. *p < 0.05, **p < 0.01 by Mann-Whitney U test. ns, not significant.

### The functional defects in neutrophils are specific to the androgenized kidney

To further specify whether androgen-dependent defects in neutrophil maturity and function are kidney specific during UTI, we performed flow cytometry with leukocyte markers on bladder tissue preparations from vehicle-treated and androgenized mice 1 and 5 dpi. Androgenized mice harbored significantly more neutrophils in the bladder 1 dpi (**Figure S5B**). By 5 dpi, the number of neutrophils in the bladders of both vehicle-treated and androgenized mice had decreased significantly (p = 0.019 and p = 0.00067, respectively; **Figure S5B**). Because the absolute cell numbers were much lower in the bladder compared with the kidney, we could only reliably ascertain age, maturity, and degranulation of the bladder neutrophils 1 dpi. As was observed in the kidney (**Figure S2A-D**), the bladders of androgenized mice 1 dpi exhibited a significantly higher proportion of immature neutrophils, and a correspondingly lower proportion of mature neutrophils, compared to vehicle-treated mice (**Figure 8A**). Of note, absolute neutrophil numbers were higher in the androgenized bladder across subtypes (**Figure 8B**). Strikingly, however, while degranulation was significantly diminished in kidney neutrophils (all subtypes) in the androgenized mice (1 dpi, **Figure 7**), degranulation by bladder neutrophils isolated 1 dpi was independent of androgen exposure (**Figure 8C-E**). Taken together, our data indicate that though androgen signaling adversely affects neutrophil maturation at multiple sites in the urinary tract, androgen-driven decrements in neutrophil function are observed specifically in the kidney environment.

**Figure 8.**
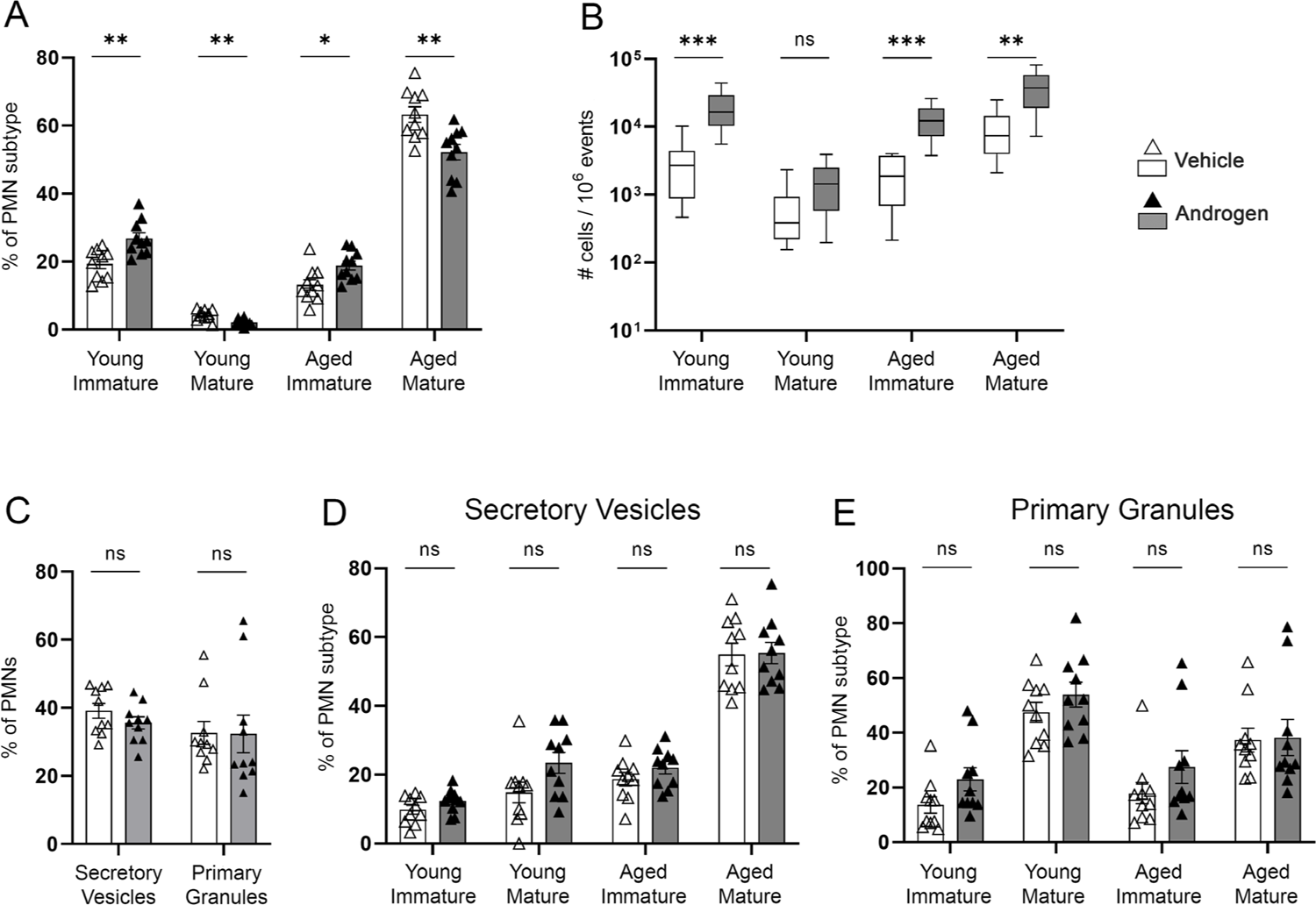
Neutrophils in the UPEC-infected bladder do not share the functional deficits seen in the kidney. (A) Distribution of neutrophil age and maturity subtypes in the bladders of vehicle-treated (open triangles, white bars) or androgenized mice (filled triangles, gray bars) 1 dpi as a proportion of total neutrophils. (B) Absolute count of each neutrophil age and maturity subtype per million events in the bladder 1 dpi. (C) Proportion of bladder neutrophils exhibiting degranulation activity 1 dpi. (D-E) Proportion of bladder neutrophils from each age and maturity subtype that had released secretory vesicles (D) or primary granules (E) 1 dpi. Each symbol represents a single mouse; n = 10 per group. **p < 0.01, ***p < 0.001 by Mann-Whitney U test. ns, not significant.

## Discussion

Urinary tract infections are very common in humans, particularly in women (55–58). While UTI in males is less frequent, it is more complicated and severe, and can exert long-term negative impacts on renal function (1,7–15). Androgen-exposed C3H/HeN mice infected with UPEC develop chronic high-titer pyelonephritis, most often accompanied by renal abscess formation. These abscesses are characterized by a large population of neutrophils surrounding tubules whose lumina harbor UPEC communities (KBCs) (16,21). The kidneys of these mice are subject to an unabating influx of CD45+ cells, predominantly reflecting steady recruitment of Ly6G+ neutrophils, but show no decreases in bacterial load at any measured time point. We therefore sought to determine how and why these recruited neutrophils in androgenized mice fail to effectively control UPEC infection. In this study, we determined that this sex difference in control of renal infection reflects an expanded population of aged, immature neutrophils in both the bladder and kidneys, and a kidney-specific decrement in neutrophil antimicrobial functions in the androgen-exposed host.

Although males have more circulating myeloid cells (neutrophils, macrophages and monocytes) than females, testosterone is recognized to exert immunosuppressive effects (23,28). Males exhibit reduced production of pro-inflammatory and increased production of immunosuppressive mediators (24–26). In our initial experiments we observed similar phenotypes, with androgenized mice harboring more CD45+ cells than vehicle-treated controls in the blood and kidneys prior to initiation of UTI. Pro-inflammatory cytokine responses in the kidneys of androgenized mice were not apparent until 7 dpi, at which point renal abscesses have already been nucleated (21). Of note, our cytokine data were obtained from whole-kidney protein extracts, possibly insensitive to cytokine production that may be very localized near intrarenal foci of UPEC at earlier time points in infection (such as in the kidneys of vehicle-treated mice). The degranulation response of neutrophils (considered in aggregate) in the kidneys of androgenized mice was similarly blunted and delayed, with release of secretory vesicles and primary granules significantly reduced compared to vehicle-treated controls until 10 dpi.

Interestingly, while the bladders of androgenized mice harbored an increased proportion of aged immature neutrophils 1 dpi compared to vehicle-treated mice, they were not inhibited in their degranulation capacity at the same time point. Taken together, these data indicate that testosterone exerts suppressive effects on the functional ability of recruited neutrophils specifically within the kidney environment, therefore hindering the androgenized host from being able to effectively resolve UTI.

Aging and maturation of neutrophils influence their ability to access sites of infection, along with their degranulation and phagocytic activity. In other model systems, aged neutrophils are more responsive to infection than young neutrophils, produce increased IL-6, and have greater phagocytic capacity (46). Meanwhile, immature neutrophils are immunostimulatory but have reduced granularity and phagocytic function compared to their mature counterparts (41,42). Although these states have been investigated separately, prior studies have not examined the combined and orthogonal effects of both age and maturation. Here, in vehicle-treated mice, aging and maturation were correlated, but we found that androgen exposure decoupled neutrophil aging from maturation within the infected urinary tract. Specifically, there was a large population of aged immature neutrophils in the kidneys of androgenized mice, comprising up to 40% of the total neutrophil population over the course of UTI. These neutrophils may have entered the bladder and kidneys as young immature cells, subsequently aging within the tissue milieu but not receiving, or properly responding to, maturation signaling. The *in vivo* maturation defect was recapitulated in C57BL/6 mice, though there were fewer aged kidney neutrophils in all tested genotypes within this background. Neutrophil maturation was restored in LysM-Cre × AR^f/f^ C57BL/6 mice, but not in Ksp-Cre × AR^f/f^ mice, indicating that androgen-dependent maturation impairment predominantly reflects myeloid cell-intrinsic AR signaling, rather than extrinsic alterations in signaling arising from androgenized kidney tissue. Of note, restoring maturation in LysM-Cre × AR^f/f^ mice did not alter kidney bacterial burden 7 dpi, indicating that androgen-dependent susceptibility to chronic, high-titer pyelonephritis is not exclusively attributable to myeloid cell AR signaling. For example, previous studies have shown that androgens influence UTI susceptibility through priming renal epithelial cells for an aberrant wound healing response, induction of pro-fibrotic macrophage polarization, and a reduction in IL-17 producing γδ T-cells (17,22,51). AR-independent effects on UTI susceptibility and outcomes are also possible. In total, our results suggest that testosterone-dependent susceptibility to severe pyelonephritis likely reflects an array of effects on multiple cell types.

The maturation defect in the kidneys of androgen-exposed mice is not the only reason for their failure to control infection despite robust recruitment, as neutrophils that have arrived specifically in the kidney exhibit diminished antimicrobial functions. Kidney and bladder neutrophils in vehicle-treated mice mustered a robust degranulation response 1 dpi, and the degranulation response in the kidney was resolved by 10 dpi, consistent with UPEC clearance in these mice. Meanwhile, only kidney neutrophils in androgenized mice exhibited dampened secretory vesicle secretion and reduced phagocytic capacity. In an apparent effort to get infection under control in the androgen-exposed host, mature neutrophils released tertiary and primary granules 10 dpi, but by this time KBCs and foci of abscess have been established for several days (21). These data indicate that while androgenization affects the maturation ability of tissue-infiltrated neutrophils, the kidney-specific environment in these mice has a compounding effect by further reducing neutrophil antimicrobial ability. Notably, though aged immature neutrophils reach UPEC within the tubular lumen, and therefore experience the same local milieu (e.g., cytokines) as other subtypes, these aged immature cells offered minimal primary degranulation, and contributed little to phagocytosis within the KBC. Interestingly, reduced neutrophil phagocytic capacity in androgenized mice was observed only in neutrophils isolated from the kidneys, reinforcing our hypothesis of a kidney-specific effect on neutrophil function. All neutrophil age/maturity subtypes isolated from kidneys of naïve androgenized mice had lower phagocytic activity compared to those of vehicle-treated mice, and this effect was even more pronounced in neutrophils isolated from kidneys 7 dpi, indicating an androgen × infection interaction on their function in the kidney.

Sex differences occur in a variety of different infections and diseases, impacting susceptibility, immunity, and responses to treatment (23,56,59). We previously demonstrated that testosterone exposure enables chronic, high-titer pyelonephritis and renal abscess formation (16,21). We now identify the cellular basis for this observation, namely a tissue environment-specific effect of androgens on neutrophil maturation accompanied by a decrement in neutrophil function that is unique to the kidney. As our model infections focused on the urinary tract, and analyses included blood, bone marrow, and spleen, it is intriguing to wonder whether other end organs, during bacterial infection, might also demonstrate analogous sex-discrepant effects on neutrophil maturation.

## Methods

### Bacterial strains

The clinical isolate of uropathogenic *Escherichia coli* (UPEC), UTI89 (60), was grown statically at 37 °C in Luria-Bertani broth (LB; Becton Dickinson, Sparks, MD). Overnight cultures were centrifuged at 7500 × *g* at 4 °C, and the resulting pellet was resuspended in sterile phosphate-buffered saline (PBS) to a final density of ∼4 × 10^8^ colony-forming units (CFU)/mL. UTI was initiated in the morning by transurethral inoculation of 50 µL of prepared bacterial suspension, delivering an inoculum of 1-2 × 10^7^ CFU. For *ex vivo* experiments, the chromosomally GFP-expressing strain UTI89 *att*HK022::COM-GFP was used (61).

### Animals

All animals were group housed in temperature-controlled suites under timed light cycles. They were supplied standard mouse chow and water ad libitum. A minimum of 2 mice (and not exceeding 5) were kept in cages with bedding and Nestlets. Experiments were conducted in female C3H/HeN mice (#040, Envigo, Indianapolis, IN; RRID:MGI:2160972). Mice were androgenized as described previously (51) via weekly intramuscular injection of 150 mg/kg testosterone cypionate (Depo-Testosterone, Pfizer, New York, NY) beginning at 5 weeks of age and continuing until sacrifice; control animals were similarly injected with cottonseed oil.

Background strains for androgen receptor knockout were originally purchased from Jackson Laboratories (Bar Harbor, ME). Ksp-Cre × AR^f/f^ mice were generated by crossing B6.129S1-*Ar^tm2.1Reb^*/J (#018450) mice with B6.Cg-Tg(Cdh16-cre)91Igr/J (#012237) to create mice homozygous for floxed androgen receptor and hemizygous for the *cre* recombinase gene under control of the cadherin 16 (kidney-specific cadherin [Ksp]) promoter. LysM-Cre × AR^f/f^ mice were similarly generated by crossing the same homozygous floxed androgen receptor mice with C57BL/6 mice expressing *cre* under the LysM promoter (kind gift from S.C. Morley). For the experiments using these strains, mice for the vehicle-treated and androgenized Cre^-^AR^f/f^ control groups were randomly chosen littermates from both Ksp-Cre × AR^f/f^ and LysM-Cre × AR^f/f^ breeders.

### Determination of bacterial loads

At the indicated time points, mice were anesthetized approximately 1 h into their light cycle with inhaled isoflurane (Patterson Veterinary, Loveland, CO), and terminally perfused with 4 °C PBS through the left ventricle. Bladders and kidneys were aseptically removed and homogenized into sterile PBS before serial dilution and plating on LB agar.

### Tissue preparation and immunofluorescence

Mice were sacrificed as described above, and harvested kidneys were placed in 4% paraformaldehyde in PBS at 4 °C for 1 h before being transferred to sterile 30% sucrose overnight at 4 °C. Kidneys were embedded and frozen in OCT (Fisher Scientific, Hampton, NH); blocks were then cryosectioned into 5-8 µm sections and mounted on Superfrost Plus slides (Fisher Scientific).

For immunofluorescence staining, sections were rinsed with PBS to remove OCT, permeabilized with 0.25% Triton X-100 (Sigma) in PBS for 10 min, blocked for 1 h with 10% fetal bovine serum (FBS; Gibco) in PBS, then stained with the following primary antibodies in blocking buffer: Goat anti-*E. coli*, O and K serotypes (1:200, Meridian Life Science #B65109G), Rat anti-Ly6G-APC (1:200, Biolegend #127614), Rat anti-CD101-PE (1:150, eBioscience #12-1011-82), Rat anti-CD49d-Alexa Fluor 488 (1:150, Biolegend #103611). Slides were washed three times with PBS before staining with Donkey anti-goat-AlexaFluor 594 secondary antibody (1:200, abcam #ab150132). Stained slides were washed again, and mounted with Prolong Gold Antifade Reagent (ThermoFisher Scientific #P36930). Images were captured with a Zeiss LSM 880 Airyscan confocal microscope (Oberkochen, Germany).

### Flow cytometry

Mice were sacrificed as described above, and harvested kidneys were manually homogenized through a 70-µm cell strainer into cold RPMI (Gibco), then centrifuged for 5 min (500 *× g*) at 4 °C. The resulting pellets were resuspended in room-temperature RBC lysis buffer (155 mM NH4Cl, 10 mM KHCO3), washed with cold FACS buffer (10% FBS, 1% w/v sodium azide in PBS), and subjected to a Percoll gradient to enrich for leukocytes. For the gradient, cell pellets were resuspended in a solution containing 36% v/v Percoll PLUS (Cytivia, Uppsala, Sweden), 25 mM sucrose in PBS, and layered on top of a solution containing 72% v/v Percoll PLUS, 25 mM sucrose in FACS buffer. Gradients were centrifuged (500 *× g*) for 30 min at 4 °C, and enriched leukocytes were collected from the buffy coat. For peripheral blood analysis, blood was collected into K2 EDTA collection tubes (BD Vacutainer #366643) via cardiac puncture before perfusion. Bone marrow was isolated from femurs. Peripheral blood, spleen, and bone marrow leukocytes were subjected to RBC lysis but did not undergo Percoll separation. Bladders were quadrisected and washed gently three times in sterile PBS to remove leukocytes from the urinary space. Washed bladders were incubated for 1 h at 37 °C in 0.34 U/mL of Liberase (Roche) in PBS. Digestion was halted by addition of FACS buffer, and the digested bladders were passed through a 70-µm cell strainer before staining. Bladder cell preparations were not subjected to hypotonic lysis or Percoll separation. Kidney, spleen, bone marrow, peripheral blood, and bladder leukocytes were stained with Live/Dead Fixable Blue (ThermoFisher Scientific) in PBS, washed, and blocked with Fc Block (BD Biosciences, San José, CA) before staining with the following extracellular antibodies in FACS buffer: MHCII-BUV395 (1:400, BD Optibuild #743876), CD19-BUV661 (1:200, BD Horizon #612971), CD45-BV510 (1:400, BD Pharmingen #563891), CD11b-BV570 (1:400, Biolegend #101233), CD18-BV650 (1:200, BD Optibuild #744600), CD49d-BV711 (1:50, BD Optibuild #740661), CD11c-BV785 (1:200, Biolegend #117336), CD3ε-SparkBlue 550 (1:200, Biolegend #100260), F4/80-BB700 (1:200, BD Optibuild #746070), CD49b-PE/Dazzle 594 (1:50, for C3H/HeN mice, Biolegend #108924), NK1.1-PE/Dazzle 594 (1:400, for C57BL/6 mice, Biolegend #108748), Ly6C-PE/Cy5.5 (1:200, Novus Biologicals #NB100-65413PECY55), CD101-AlexaFluor 647 (1:100, BD Pharmingen #564473), Ly6G-AlexaFluor 700 (1:800, Biolegend #127621), CD63-APC/Cy7 (1:200, Biolegend #143908), CXCR4-BUV805 (1:200, BD Optibuild #741979), CD62L-PerCP (1:100, Biolegend #104430), CXCR2-PE/Cy7 (1:200, Biolegend #149316). Cells were washed, resuspended in FACS buffer, and analyzed with an Aurora flow cytometer (Cytek Biosciences, Fremont, CA) and FlowJo software (BD Biosciences). A representative gating scheme is provided in **Figure S6**.

### Ex vivo phagocytosis assay

Kidneys were harvested and prepared as described in the preceding section. After the Percoll gradient, 1 × 10^6^ cells (by hemacytometer) were incubated with 1 × 10^6^ CFU of UTI89 *att*HK022::COM-GFP (Wright et al., 2005) in RPMI 1640 (Gibco) + 10% FBS for 30 min at 37 °C with 5% CO2. Cells were then washed with FACS buffer, and staining was performed as described in the preceding section. The gating of GFP+ UPEC is shown in **Figure S6**.

### Protein extraction

Harvested kidneys were flash frozen in liquid nitrogen and stored at −80 °C until use. Kidneys were homogenized in RIPA buffer (50 mM Tris-HCl, 150 mM NaCl, 1% v/v Nonidet P-40, 0.1% w/v SDS, 0.5% w/v sodium deoxycholate, pH 7.4) containing PhosSTOP phosphatase inhibitor (Roche; Basel, Switzerland) and cOmplete Mini protease inhibitor (Roche). Lysates were cleared by centrifugation (2 × 5 min in a tabletop microcentrifuge at maximum speed), and protein concentration was determined by BCA assay (Invitrogen, Carlsbad, CA).

### Cytokine quantification

Extracted whole-kidney protein was diluted in PBS to 900 µg/mL and subjected to a Bio-Plex Mouse Cytokine 23-plex Assay (#M60009RDPD, Bio-Rad, Hercules, CA), according to the manufacturer instructions. The plate was read with a Bio-Plex 200 system and analyzed using BioPlex Manager 6.1 software.

### Statistical analysis

Statistical analyses were performed by unpaired, non-parametric Mann-Whitney U-tests using GraphPad Prism 9.5.0. P-values <0.05 were considered significant. Statistical details for each figure can be found in the corresponding figure legend.

### Study approval

All animal protocols received prior approval from the Washington University Institutional Animal Care and Use Committee.

## Author contributions

Conceptualization, T.N.H. and D.A.H.; Methodology, T.N.H. and C.A.C.; Investigation, T.N.H, C.A.C, and D.A.H.; Writing – Original Draft, T.N.H. and D.A.H.; Writing – Review & Editing, T.N.H., C.A.C., and D.A.H.; Visualization, T.N.H. and D.A.H.; Funding Acquisition, D.A.H.; Supervision, D.A.H.

## Acknowledgments

This work was supported by grants from the National Institutes of Health (R01-DK111541, R01-DK126697, and R01-AI158418 to D.A.H.) and from the Children’s Discovery Institute through the Washington University Center for Cellular Imaging (CDI-CORE-2019-813). T.N.H. was supported by NIH grant T32-DK007126. We thank Dr. Gina Clemens for advice and discussions regarding neutrophil assays.

