## Supplemental Figures for "Androgen exposure impairs neutrophil maturation and function within the infected kidney"

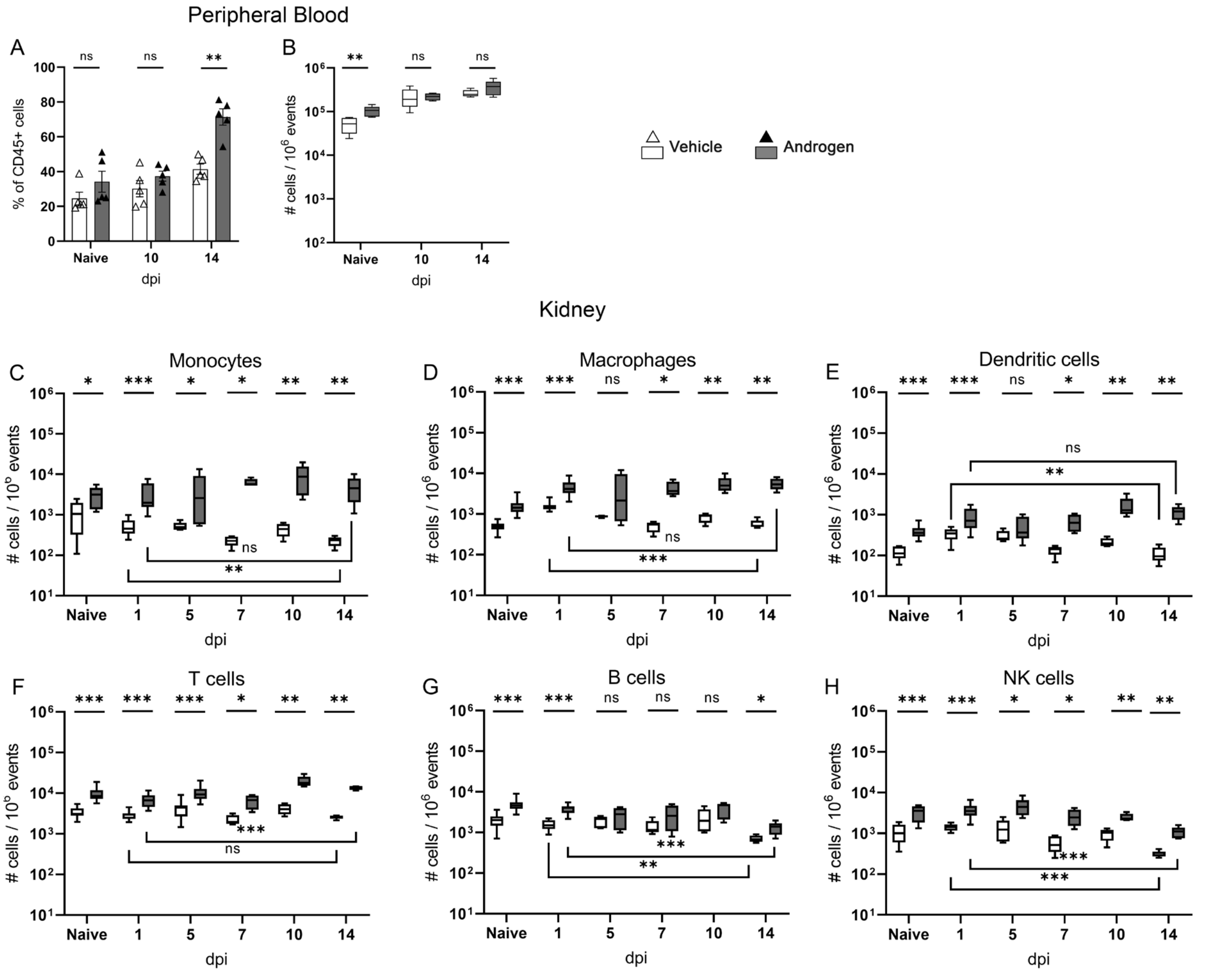


**Figure S1.** Timeline of the neutrophil population in peripheral blood and leukocyte populations in the kidneys. The population of neutrophils in the peripheral blood, as a proportion of CD45+ cells (A) and absolute count (per million events; B) are shown for vehicle-treated (open triangles, white bars) and androgenized mice (closed triangles, gray bars). (C-H) Leukocyte subset populations over time in the kidneys of vehicle-treated (white bars) and androgenized mice (gray bars). Bars indicate mean with SEM, Box and whisker plots represent 95 percentile range with min and max. Each symbol represents a single mouse; n = 5-15 per time point. *$p < 0.05$, **$p < 0.01$, ***$p < 0.001$ by Mann-Whitney U test. ns, not significant.

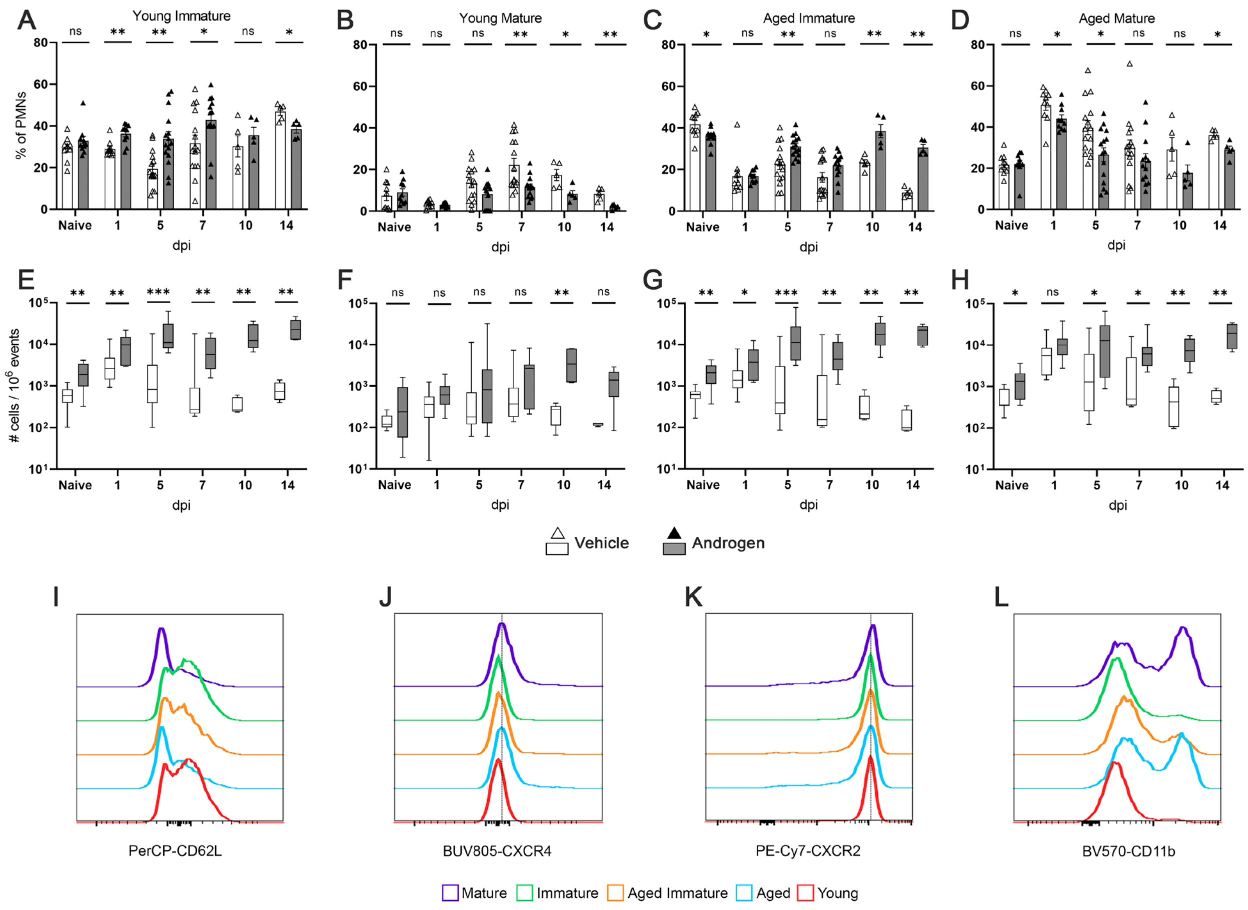


**Figure S2.** Populations of neutrophil age and maturity subtypes in the kidneys throughout infection. Relative proportion of all neutrophils (PMNs; A-D) and absolute number of neutrophils (per million events; E-H), as measured by flow cytometry, represented by young immature (A, E), young mature (B, F), aged immature (C, G), or aged mature (D, H) subtypes, in the kidneys of vehicle-treated (open triangles, white bars) or androgenized mice (closed triangles, gray bars). (I-L) Histograms of expression of CD62L (I), CXCR4 (J), CXCR2 (K) and CD11b (L) in a representative sample of neutrophils gated solely on maturity (purple; mature, green; immature) or age (blue; aged, red; young), compared to aged immature neutrophils (orange). Bars indicate mean with SEM. Box and whisker plots represent 95 percentile range with min and max. Each symbol represents a single mouse; n = 5-15 mice per time point. *$p < 0.05$, **$p < 0.01$ by Mann-Whitney U test. ns, not significant.

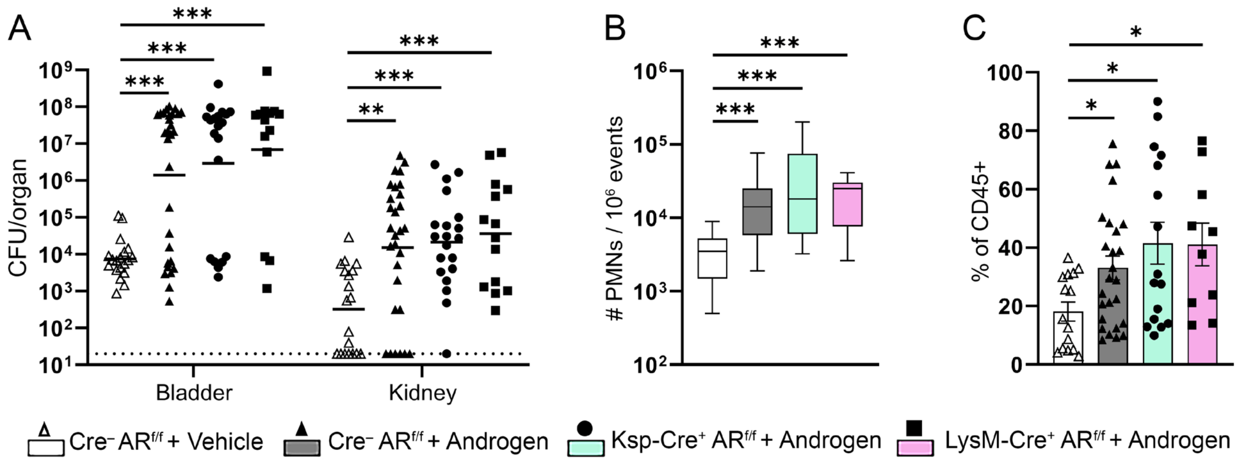


**Figure S3.** CFU and neutrophil recruitment in conditional AR-deficient C57BL/6 strains. Androgenized C57BL/6 mice exhibit chronic high-titer pyelonephritis, even with conditional AR deficiencies. (A) Bladder and kidney bacterial loads in vehicle-treated $Cre^{-}AR^{f/f}$ (open triangles, white bars), androgenized $Cre^{-}AR^{f/f}$ (closed triangles, gray bars), androgenized Ksp-Cre × $AR^{f/f}$ (squares, green bars), or androgenized LysM-Cre × $AR^{f/f}$ (circles, pink bars) mice 7 days post infection. Lines indicate geometric mean. (B) Kidney neutrophil (PMN) counts in the same groups of mice (per million events). Box and whisker plots represent 95 percentile range with min and max. (C) Kidney neutrophils in the same groups of mice as a percentage of CD45+ cells. Bars indicate mean with SEM. n = 15-26 mice per group; $*p < 0.05$, $**p < 0.01$, $***p < 0.001$ by Mann-Whitney U test. CFU, colony-forming units.

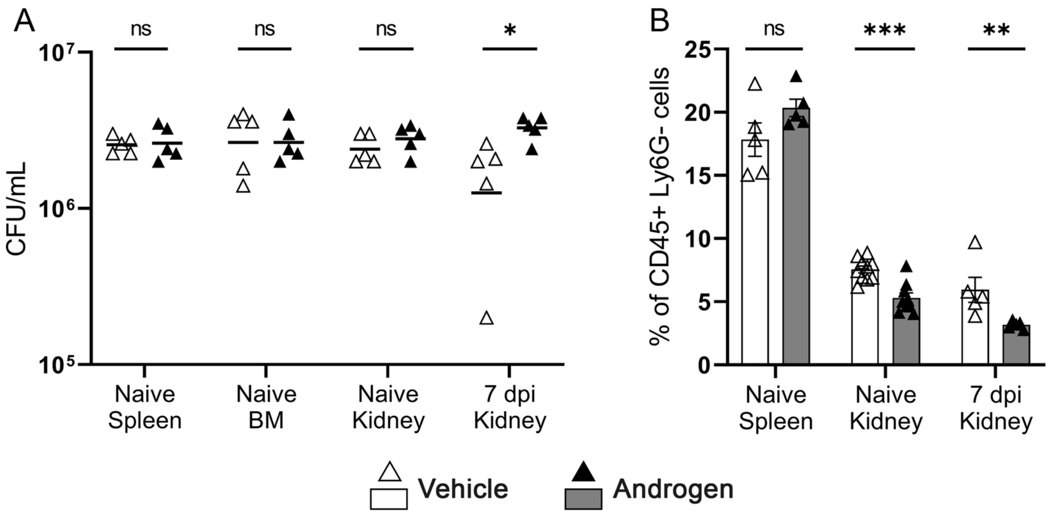


**Figure S4.** Remaining UPEC after ex vivo phagocytosis assay and phagocytic capacity of non-neutrophil leukocytes in C3H/HeN kidneys. (A) Bacterial colony-forming units (CFU) of GFP+ UPEC in supernatants after 30 min of exposure to non-neutrophil leukocytes (CD45+ Ly6G–) isolated from naïve spleen, bone marrow (BM), or kidneys, or from kidneys 7 days post infection (dpi) with UPEC, in vehicle-treated (open triangles) or androgenized mice (closed triangles). Lines indicate geometric mean. (B) Phagocytic capacity (as measured by % GFP positive) of CD45+ Ly6G– leukocytes isolated from naïve spleen or kidneys, or from kidneys 7 dpi, of mice treated with vehicle (white bars) or androgen (gray bars). Bars indicate mean with SEM. n = 5 mice per group; $^{**}p < 0.01$, $^{***}p < 0.001$ by Mann-Whitney U test. ns, not significant.

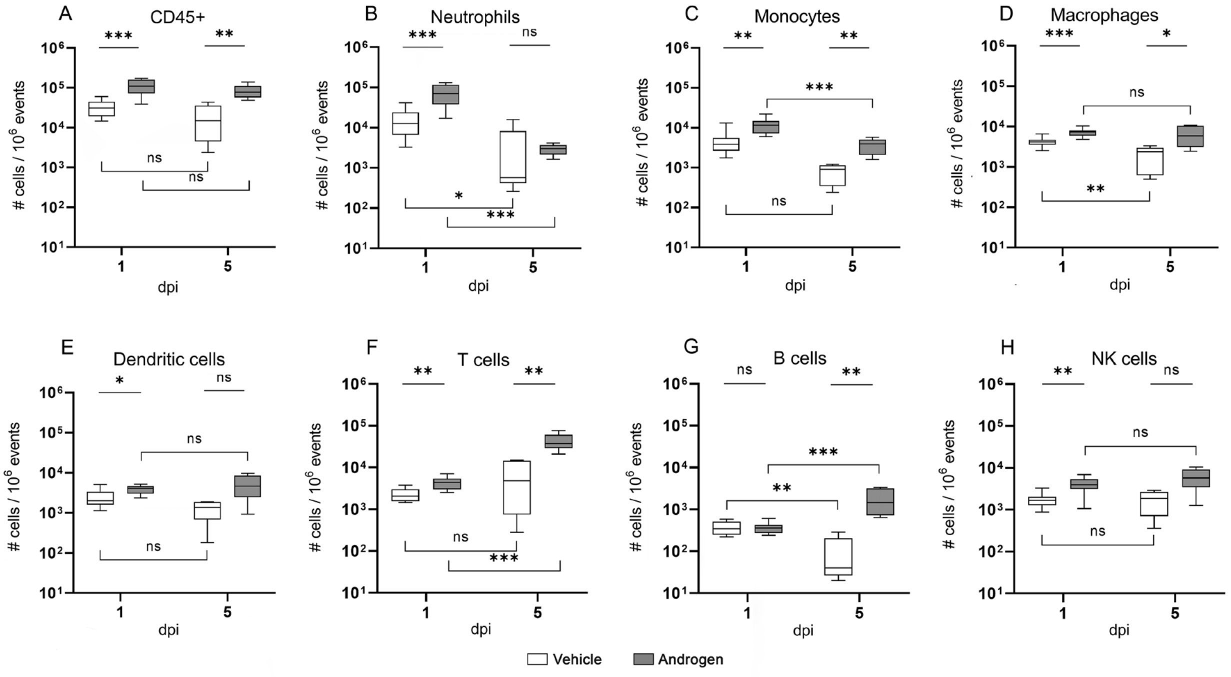


**Figure S5.** Timeline of leukocyte populations in the bladder. (A-H) Absolute count per million events, in vehicle-treated (white boxes) and androgenized mice (gray boxes), of all leukocytes (A) and leukocyte subset populations (B-H) in the bladder 1 and 5 dpi. Box and whisker plots represent 95 percentile range with min and max. n = 5-10 per time point. $*p < 0.05$, $**p < 0.01$, $***p < 0.001$ by Mann-Whitney U test. ns, not significant.

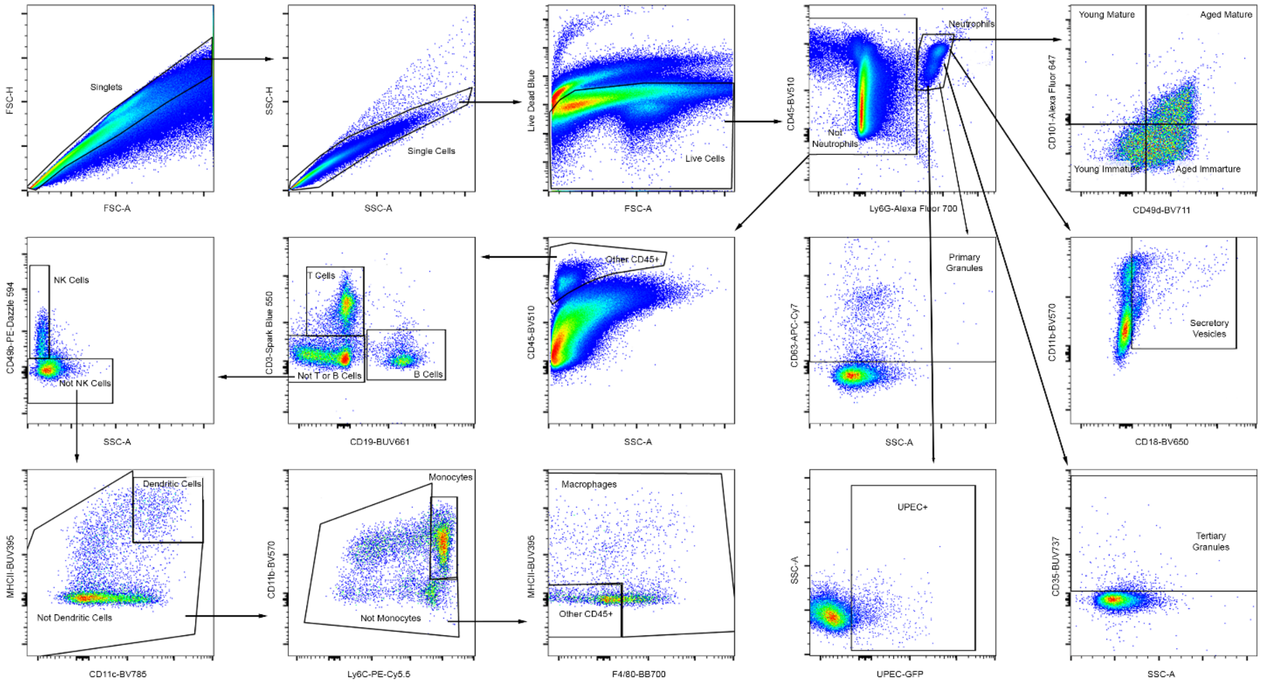


**Figure S6.** Representative gating scheme for analyzing leukocytes and neutrophils from mouse kidneys.
